# scCODE: an R package for personalized differentially expressed gene detection on single-cell RNA-sequencing data

**DOI:** 10.1101/2021.11.18.469072

**Authors:** Jiawei Zou, Miaochen Wang, Zhen Zhang, Zheqi Liu, Xiaobin Zhang, Rong Hua, Ke Chen, Xin Zou, Jie Hao

**Affiliations:** School of Life Sciences and Biotechnology, Shanghai Centre for Systems Biomedicine, Shanghai Jiao Tong University, Shanghai, China; Institute of Clinical Science, Zhongshan Hospital, Fudan University, Shanghai, China; Jinshan Hospital Center for Tumor Diagnosis & Therapy, Jinshan Hospital, Fudan University, Shanghai, 201508, China; Department of Oral and Maxillofacial-Head & Neck Oncology, Shanghai Ninth People’s Hospital, Shanghai Jiao Tong University School of Medicine; College of Stomatology, Shanghai Jiao Tong University; National Center for Stomatology; National Clinical Research Center for Oral Diseases; Shanghai Key Laboratory of Stomatology; Shanghai Key Laboratory of Plant Functional Genomics and Resources, Shanghai Chenshan Botanical Garden, Shanghai, 201602, China; Department of Thoracic Surgery, Shanghai Chest Hospital, Shanghai Jiao Tong University, Shanghai, China; Department of Cardiovascular Surgery, Shanghai Chest Hospital, Shanghai Jiao Tong University, Shanghai, China

**Keywords:** bioinformatics, scRNA-seq data, differentially expressed gene detection

## Abstract

Differential expression (DE) gene detection in single-cell RNA-seq (scRNA-seq) data is a key step to understand the biological question investigated. We find that DE methods together with gene filtering have profound impact on DE gene identification, and different datasets will benefit from personalized DE gene detection strategies. Existing tools don’t take gene filtering into consideration, and couldn’t evaluate DE performance on real datasets without prior knowledge of true results. Based on two new metrics, we propose scCODE (single cell Consensus Optimization of Differentially Expressed gene detection), an R package to automatically optimize DE gene detection for each experimental scRNA-seq dataset.

## Background

In the past few years, stupendous development in single-cell RNA sequencing (scRNA-seq) has made it a powerful technology in transcriptomic biology. scRNA-seq data can provide unprecedented transcriptomic profile of individual cells. For the interpretation of scRNA-seq dataset where comparison scenarios occur, for example, cell development[1], cancer research[2, 3] and immune research[4, 5], etc., the detection of DE gene is critical for downstream researches. Accurate and reliable DE gene detection can accelerate the discovery of novel regulator genes and biomarkers[6, 7].

In order to detect the true DE genes in scRNA-seq data, various DE methods and algorithms have been developed, including methods particularly designed for scRNA-seq data, such as MAST[8], BPSC[9] and scDD[10], methods previously used in bulk data, such as edgeR[11], DESeq2[12] and the more common methods such as t-test[13] and Wilcoxon-test[13] et al. And there also exists method[14] integrating these methods.

Although there exist several tools to assess the performance of DE methods on cohort of scRNA-seq data[15, 16], they only evaluate each method based on the average performance on many scRNA-seq datasets. However, the method selected this way may not always be suitable for every dataset, there’s still no solution to optimize DE gene detection performance for a specific experimental scRNA-seq dataset. In addition, there is lack of systematic evaluation on gene filtering methods, regardless its profound influence in scRNA-seq data interpretation[17, 18].

Here, we first systematically evaluated the performance of combinations of different gene filtering and DE methods based on the metrics used in conquer[16], we exhibited the true influence of gene filtering and that different analysis strategy varies a lot in DE gene detection results. Besides, we noticed that each dataset requires specific method to achieve its optimal result, but there is lack of means to determine the most suitable method for a given dataset without prior knowledge of the true DE genes. Thus, we constructed two novel metrics, CDO (Consistent DE genes Order) and AUCC (Area under consistency curve). Principally, the two metrics rate the performance of DE gene detection strategies by evaluating their efficiency of capturing common genes. The reliability of the two metrics was justified on simulated datasets with known true DE genes, and both metrics showed significant correlation with the AUROC (Area Under Receiver Operating Characteristic Curve) which was regarded as the ground truth. Based on the new metrics, we constructed scCODE (single cell Consensus Optimization of Differentially Expressed gene detection), an R package to provide optimized DE gene results based on the consensus genes obtained by the top suitable analysis strategies. We found that scCODE demonstrated better and more robust performance, while other analysis strategies showed unstable performance on different simulated datasets. The scCODE was further justified on a mouse scRNA-seq data[19] comparing activated CD4+ T cells with naïve CD4+ T cells. The obtained biological processes associated with T cell activation highly agreed with the experimental design of the origin study[19].

## Methods

In this article, we focused on the DE gene detection of two-group comparison. First, systematic evaluation of various analysis strategies by combining gene filtering (conquer[16], OGFSC (Optimal Gene filtering for Single-cell data)[18], scmap[20] and no-filter) and DE methods was performed based on the following metrics, i.e., FPR (False positive Rate), SNR (Signal-Noise Ratio), FDP (False discovery Proportion), TPR (True Positive Rate) and AUROC (Figure. 1a). However, the above metrics are not applicable in real scenarios as they all require prior knowledge of true DE genes. To tackle this issue, two new metrics CDO and AUCC were constructed for evaluating DE gene detection performance in experimental scenarios. Using the new metrics, we designed scCODE platform to optimize the DE gene detection performances (Figure. 1b).

**Figure. 1.**
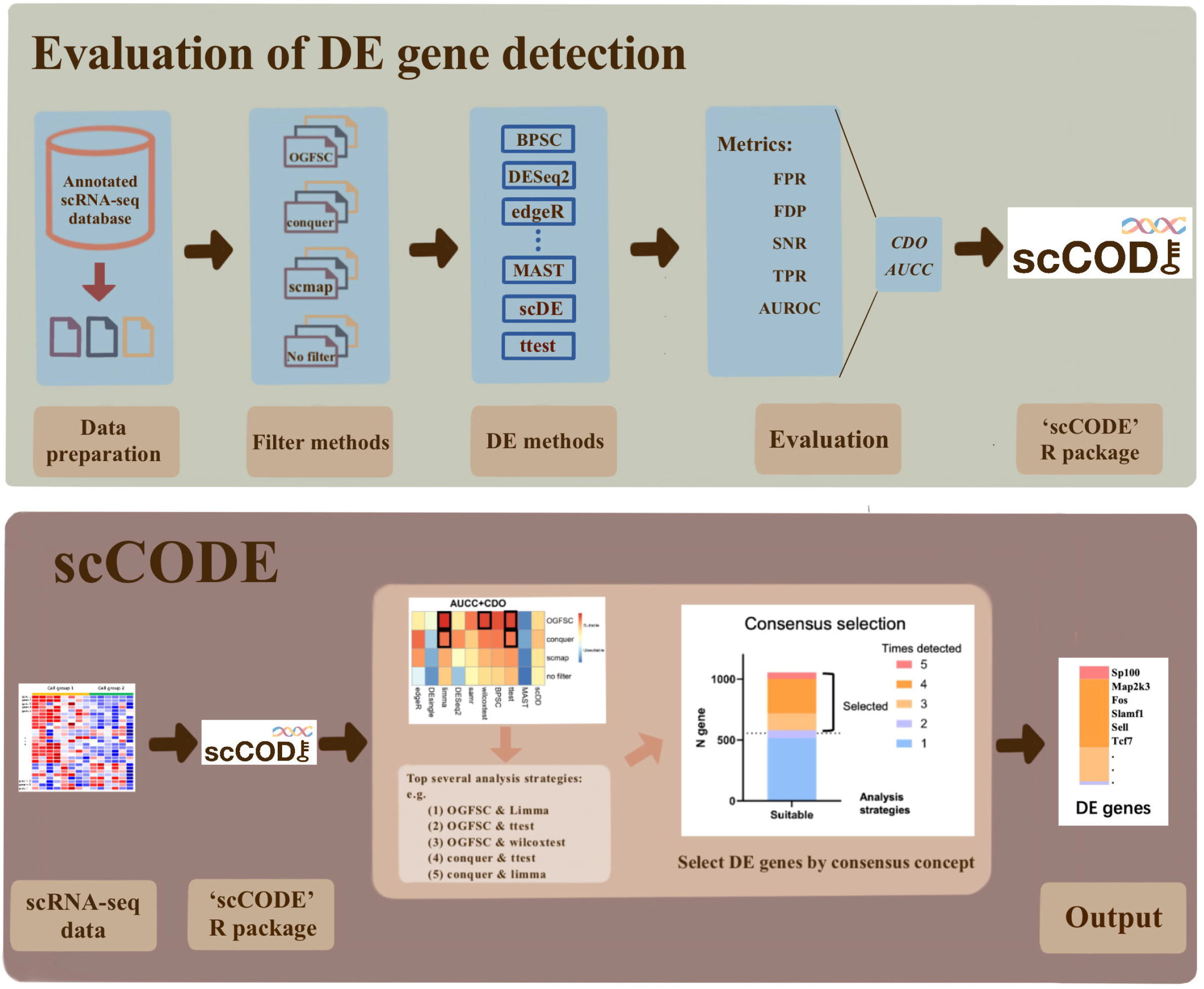
The Evaluation of DE gene detection performance and the schematic of scCODE. (a) The evaluation of the performance of different analysis strategies. (b) The schematic of scCODE.

### Single cell RNA-seq data

We collected 20 scRNA-seq datasets from the conquer[16] database, including 3 datasets of UMI protocol and 17 datasets of full-length ones. These datasets varied in organism (mouse or human), cell type, sample size and other aspects. The scRNA-seq data used in this article was categorized into 3 types: simulated data without true DE genes (null data), simulated data with true DE genes and real experimental datasets.

### Simulated data without true DE genes (null data)

We generated multiple null instances from each experimental dataset by sampling from one type of cell, where no true DE gene was expected. One type of cells from each real dataset was selected as the source data where the null data sampled from. A stratification strategy was applied on the sample size, the number of cells in each sampling varied from 20%, 40%, 60%, 80% to 100% of the total number of the cells of the source dataset, and each sampling was replicated 5 times, thus we got 25 null instances from each experimental dataset.

#### Simulated data with true DE genes

Using the principle similar as above, we first generated null simulate data. Then, all cells were repeatedly resampled into two groups, while the gene expression of one cell group was multiplied by a fold-change value. In the simulated datasets, 2% of the genes were randomly selected as DE genes, the fold change of which depended on a gamma distribution with shape 4 and base varied in 2,3,4,5,6,7,8,10.

#### Experimental data

As mentioned before, we collected 20 real datasets, some of which contain more than two types of cells. And only two types of cells were selected as origin experimental data for DE gene detection between two groups of cells.

More information is detailed in the data information table (Supplementary table 1).

### Evaluation of DE gene detection performance

#### Gene filtering methods

Although filtering genes with high-level of noise is already applied in many researches[21-23], there is no golden standard for gene filtering in scRNA-seq data. The methods used in most studies are hard-thresholding methods based on expression level, i.e., genes with expression value below threshold would be excluded. Conquer was included in this study as an option for hard thresholding. Another approach, OGFSC[18], which builds multiple candidate thresholding curves between CV^2^ (coefficient of variation) and gene expression level μ, and provides an optimal thresholding curve to preserve biological signal and reduce technical noise. We also included feature selection function from scmap[20] as an alternative gene filtering method here, which ranks the information contained in each gene. The 3 gene filtering methods were benchmarked against no-filter strategy.

##### Conquer

conquer[16] is a representational filtering method of hard thresholding, which has been widely applied in scRNA-seq data analysis. Conquer filters genes with an expression above 1 count in less than 25% of the cells. In this article, we made it filter genes with an auto-adjusted threshold (5% - 25%) in order to maintain sufficient number of genes (>=2000).

##### OGFSC

OGFSC[18] is a tool developed particularly for filtering genes in scRNA-seq data, which distinguishes signal from noise based on a soft thresholding curve of expression level and variance. In this article, the thresholding curve is automatically determined by OGFSC.

##### Scmap

The feature selection of scmap[20] is not designed particularly for filtering genes, but it’s power of sorting the informative genes is similar to that we concerned about. In this article, the function of feature selection in scmap is treated as an alternative gene filtering method, and the number of remained genes (top features) is set to 5000.

#### DE methods

There are plenty of DE methods for scRNA-seq data, here we examined 10 methods (Table 1). Some methods are usually used in bulk data, including edgeR[11], DESeq2[12] and limma[24]. For edgeR, we applied likelihood ratio test (LRT)[25] with default TMM normalization and default dispersion estimation. DESeq2 was run on rounded data by default settings. Voom-limma was applied on counts data by default settings. We also included 2 nonparametric methods, Wilcoxon test and SAMseq[26]. The Wilcoxon test were applied on log-transformed data with default settings. SAMseq was run on count data with permutations = 1000. Besides, 4 methods particularly designed for scRNA-seq data were applied, including MAST[8], BPSC[9], scDD[10] and DEsingle[27]. We applied MAST to log-transformed counts data without considering the cellular detection rate. BPSC and DEsingle were applied to count data with default settings. scDD was run on count data, with prior parameter alpha=0.01, mu0=0, s0=0.01, a0=0.01, b0=0.01. Finally, we also applied a non-paired t-test[13] on log-transformed data.

**Table 1.**
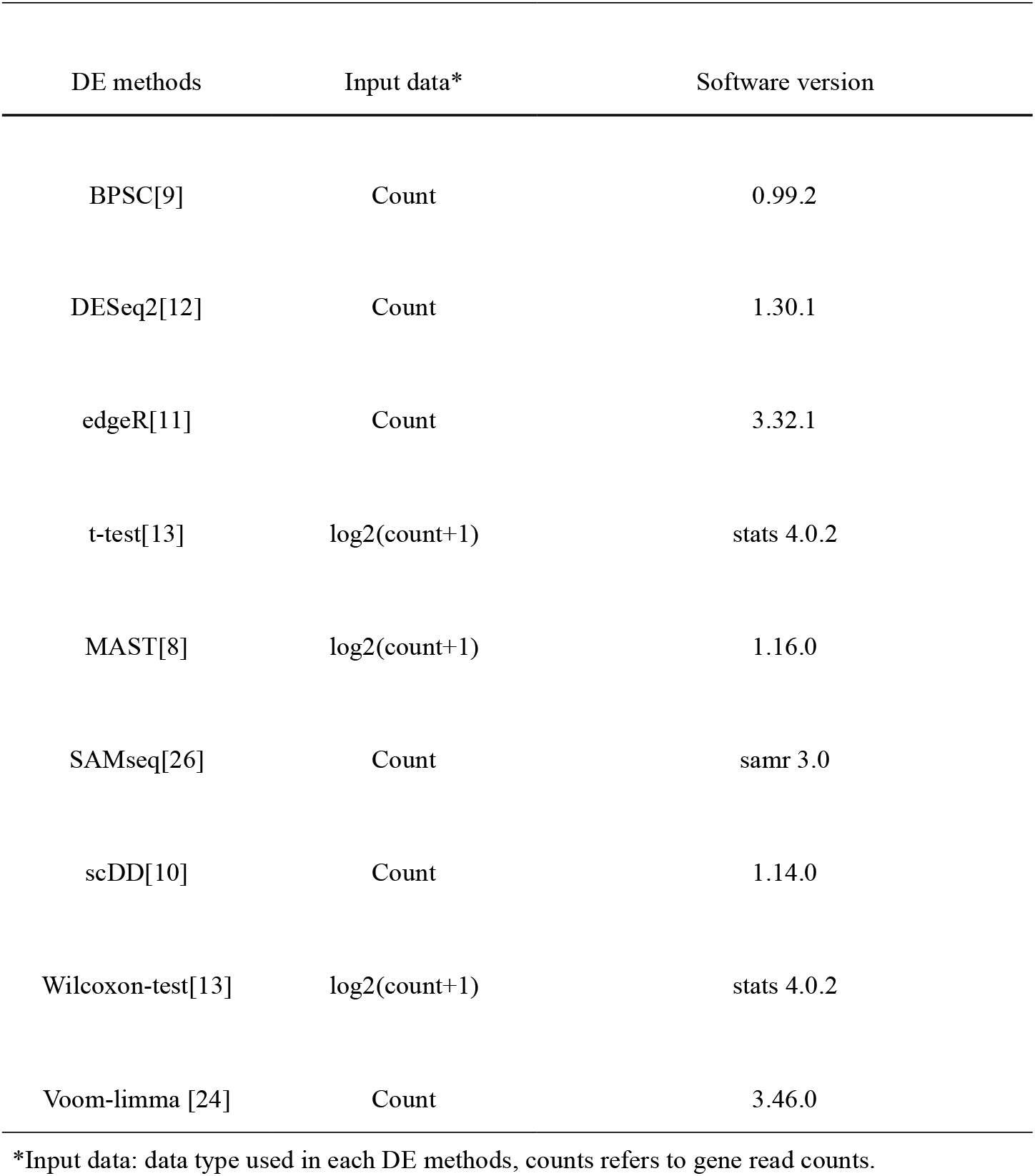
DE methods and algorithm used in this article.

**Table 2.**
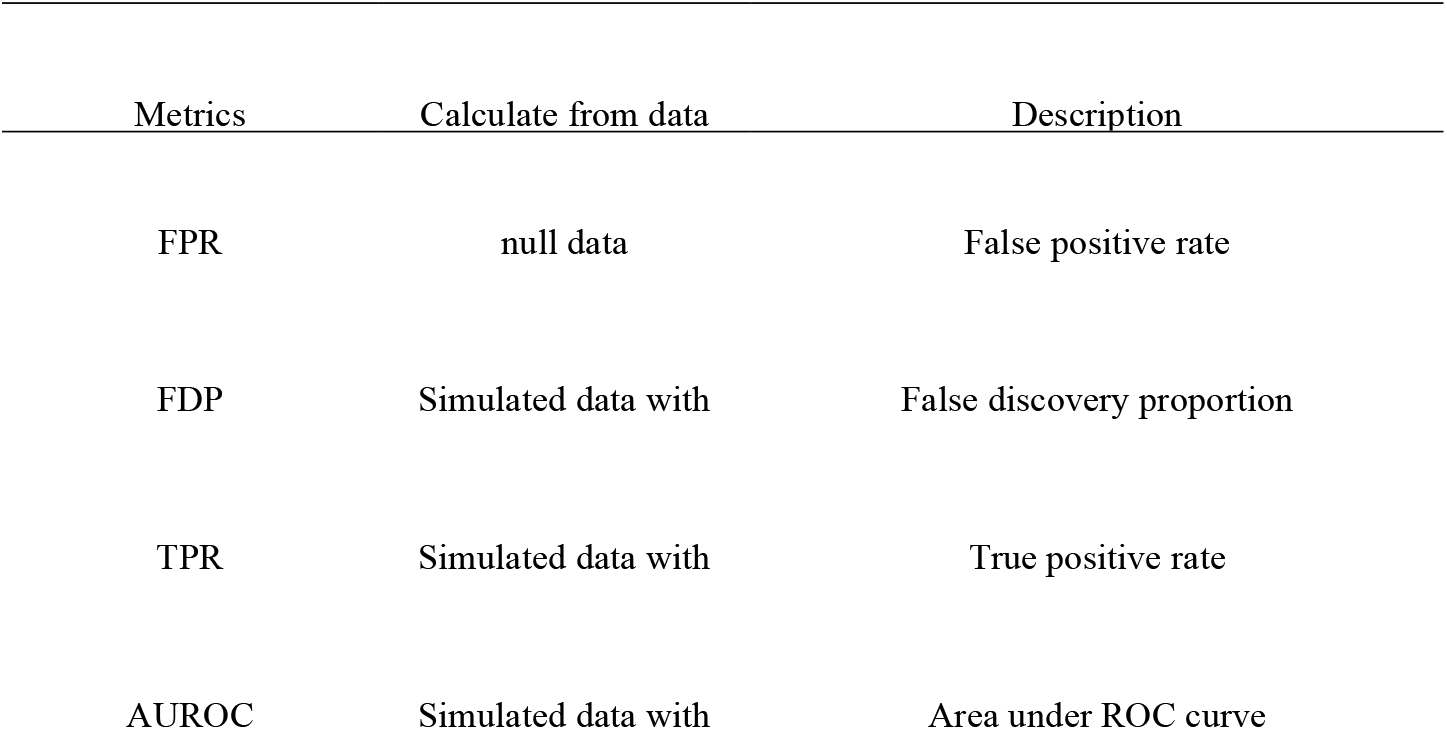
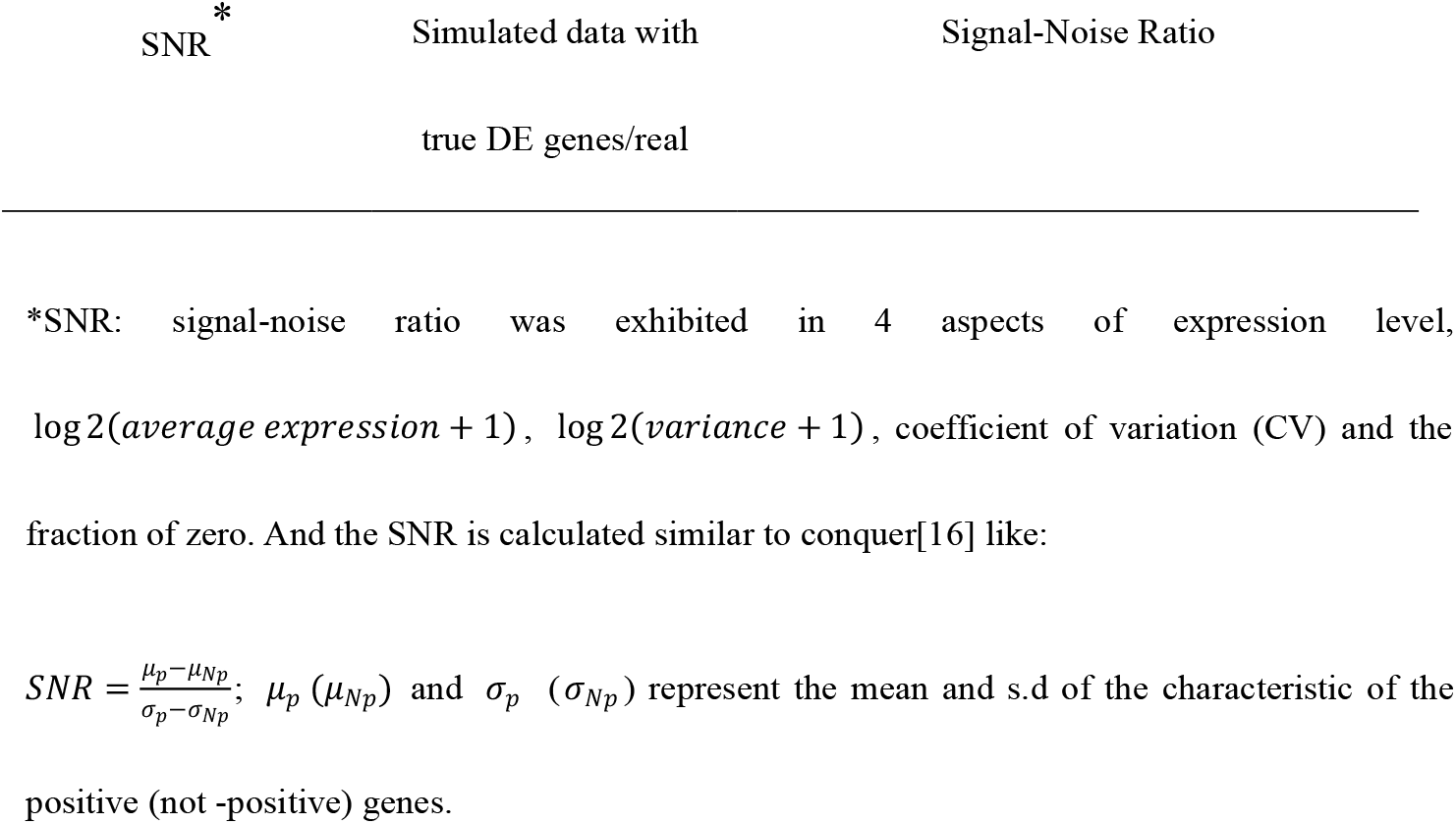
Metrics used in evaluation of analysis strategies for DE gene detection.

#### Metrics

The evaluation of the DE gene detection was based on several criteria commonly used before. FPR was calculated on null data, and genes detected with p-value ≤ 0.05 were recognized as false positive results. FDP, TPR and AUROC were calculated from simulated data with true DE genes, whereas genes with adjusted p-value (‘fdr’) ≤ 0.05 were assigned as positive DE genes. SNR was calculated on both simulated data with true DE genes and experimental data by comparing the four characteristics between positive genes detected and non-positive ones.

### scCODE

#### scCODE metrics

The above metrics are highly dependent on the prior knowledge of true DE genes, which is not always available. AUROC is the golden standard widely applied, and it’s defined as below:

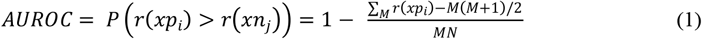

where *r*(*x*) represents the rank of instance *x, xp*_*i*_ represents a randomly selected true positive instance, *xn*_*j*_ represents a randomly selected true negative instance; *M* represents the number of true positive instance, *N* represents the number of true negative ones.

To approximate AUROC in real cases, two metrics CDO and AUCC were used here. The basic concept of CDO and AUCC is similar to AUROC but requires no prior knowledge. CDO regards the most frequently identified positive genes as seed DE genes. It further reflects the ability of each method to detect these seed genes as positive results before other genes by calculate their relative rank sorted by p-value in the DE results of each analysis strategy. AUCC is not a new metric designed by us, it’s used to calculate the consistency between two pair of results. We assume that positive DE genes detected by more methods as more reliable, and the method with a higher CDO and AUCC is also considered as more trustworthy.

##### CDO

Consistent DE gene Order, is a novel parameter proposed here, range from 0 to 1.

Basically, the common DE genes detected by no less than 75% of the different analysis strategies are considered as seed DE genes. For DE gene detection in experimental data, we consider the seed DE genes as a part of the true positive results. if we could identify *M*1 seed DE genes, we calculate CDO like below:

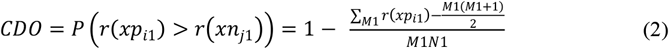

where *M*1 < *M*; *N*1 > *N*.

We find that:

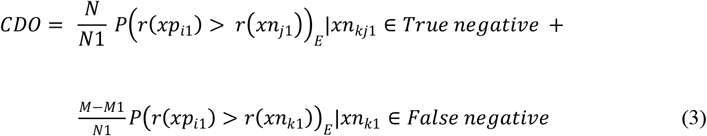

While

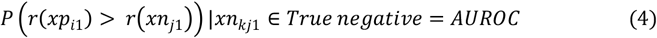

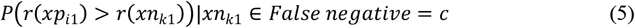

where *c* is a constant which depends on the bias of seed DE genes, and 0 ≤ *c* ≤ 1.

So

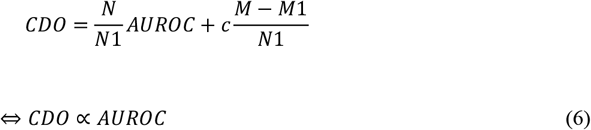

There is a liner relationship between *CDO* and *AUROC*.

In practice, we make the number of seed DE genes *M*1 no less than 5 in order to increase the robustness of seed DE genes.

##### AUCC

AUCC (Area under Concordance Curve) [16], a measurement of concordance. In this article, we calculated the AUCC value between different DE methods but with the procession of the same gene filtering method. We build concordance curve for each pair of DE methods, for top *n* genes, and the AUCC value of each DE method is the average AUCC value of itself with the other 9 DE methods. AUCC can be calculated like below:

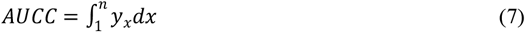

where *y*_*x*_ is the number of common results between the two strategies, *n* represents the top *n* DE gene results.

#### Consensus optimization

Using CDO and AUCC, we are able to identify optimal strategies for the specific scRNA-seq dataset. Based on the DE genes detected by various analysis strategies, scCODE produces consensus DE gene results. scCODE summarizes the top (default as all) DE genes from each of the strategy selected. The principle of consensus optimization is that the DE genes with higher frequency of observation by different analysis strategies are more reliable. We rank the genes by their number of observations. For the genes with same number of observations, they are sorted by absolute fold changes. Finally, the top n (median number of the DE genes detected from each analysis strategy) genes are used as final result from scCODE.

## Results

For evaluation of DE gene detection, we adopted 4 gene filtering approaches and 10 DE methods, which lead to 40 analysis strategies (see Methods). Each data instance was first filtered with the number of retained genes no less than 2000. From the remaining genes, DE ones were detected using various DE methods. The results of 40 analysis strategies for each data instance were systematically evaluated by the above metrics. In this article, while calculating the CDO and AUCC metrics, the top n (default as 500) DE genes detected by each analysis strategy were considered. Based on CDO and AUCC, scCODE provided personally optimized DE gene results for any given scRNA-seq dataset.

### The evaluation of FPR on null dataset without true DE genes

Using null datasets, we evaluated the type I error control power of different analysis strategies by FPR. We recorded the proportion of genes that were assigned with a p-value lower than 0.05. Similar to the evaluation in conquer[16], we found that most of these DE methods could keep the FPR near to the imposed level. Some DE methods resulted an FPR higher than the imposed level such as edgeR and limma, while other methods seemed to achieve an FPR value under the theoretical level (Figure. 2). Furthermore, the FPRs after gene filtering procedure were generally lower than those without filtering in most cases. For the DE methods that could keep the FPR value well under 0.05, such as scDD, MAST and t-test, FPR seemed to be optimized by coupling with OGFSC gene filtering. Overall, the FPR evaluation shows that gene filtering before DE gene detection does help to decrease the FPR. We also demonstrated the relationship between FPR and cell sample size. A cubic regression for each method shows that the cubic curve of FPR waved up when sample size increased (Supplementary Figure. 1). And we observed that the curves of the same DE method were very similar in shape with various filtering methods. Therefore, the relation between FPR and sample size depends more on DE methods.

**Figure. 2.**
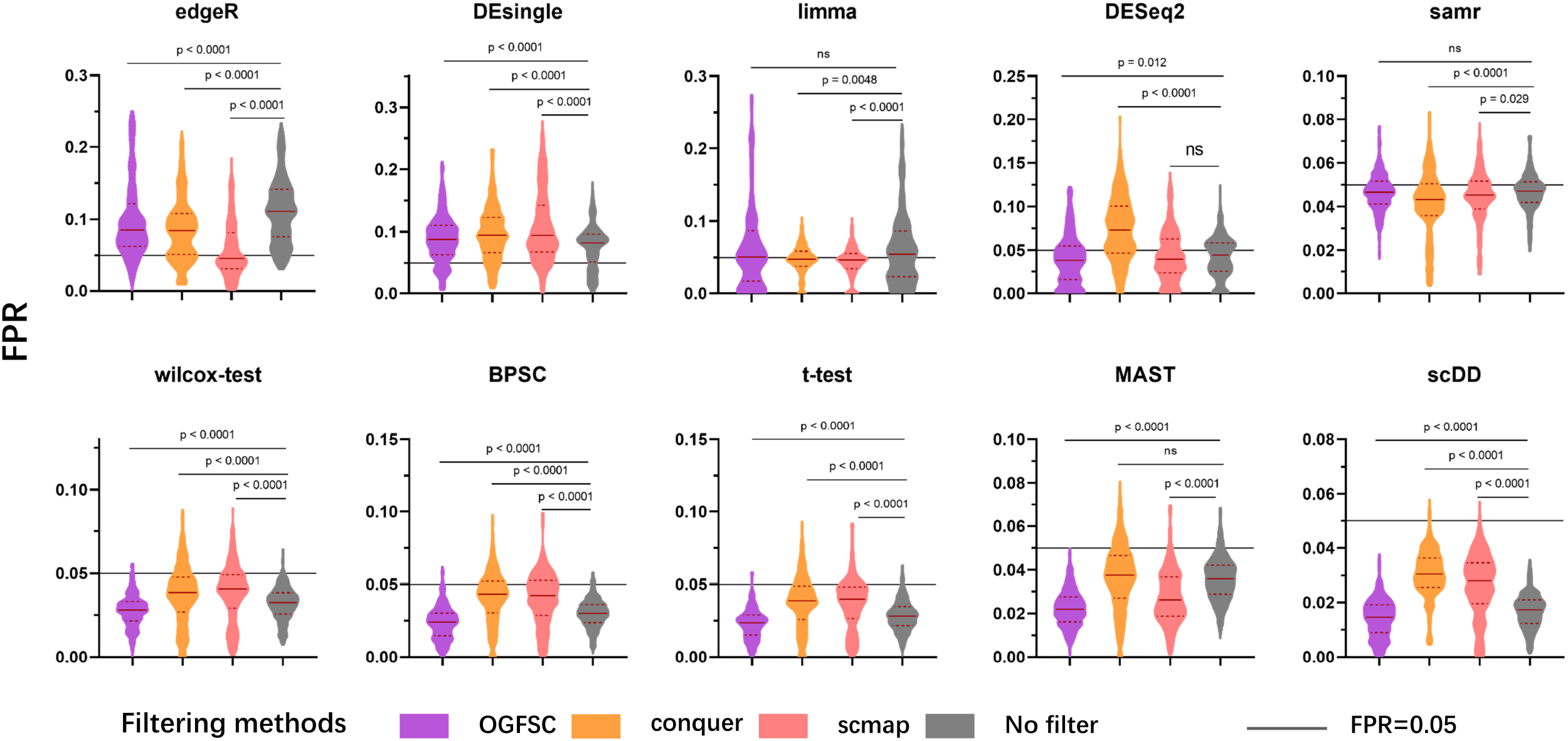
The violin plot of FPR results of different analysis strategies. The results are exhibited separately by diverse DE methods, and 4 different colors of violins represents 4 gene filtering methods. Each violin contains the FPR values resulted from every null data instance, central line, median value; dotted line, first and third quantile. Outliers (larger than third quantile + 2*IQR or smaller than first quantile – 2*IQR) are removed.

### The evaluation of SNR on data containing DE genes

In order to evaluate the DE gene detection bias, we compared the characteristics of genes detected as DE (adjusted p-value < 0.05) with that non-DE ones. The characteristics include the log 2(*average expression* + 1), log 2(*variance* + 1), coefficient of variation (CV) and the fraction of zero (the expression level) among all cells. We calculated SNR on both simulated data and experimental datasets with known true DE genes. And only the results with more than 5 genes detected as DE were recorded as valid results. In fact, the results on experimental datasets were largely similar with that of simulated data with true DE genes (Figure. 3, Supplementary Figure. 2). We noticed that the average expression level, variance and CV of the DE genes detected by either filtering methods and DE methods were significantly higher than that of non-DE ones (Figure. 3a, b, c), while the fraction of zeros was mostly lower (Figure. 3d). Furthermore, considering the performance of different filtering methods, we observed that the SNR of each characteristic after filtering by scmap and conquer was closer to 0. It indicated that the DE genes detected by any analysis strategies with scmap or conquer shared a more similar characteristic with those assigned as non-DE.

**Figure. 3.**
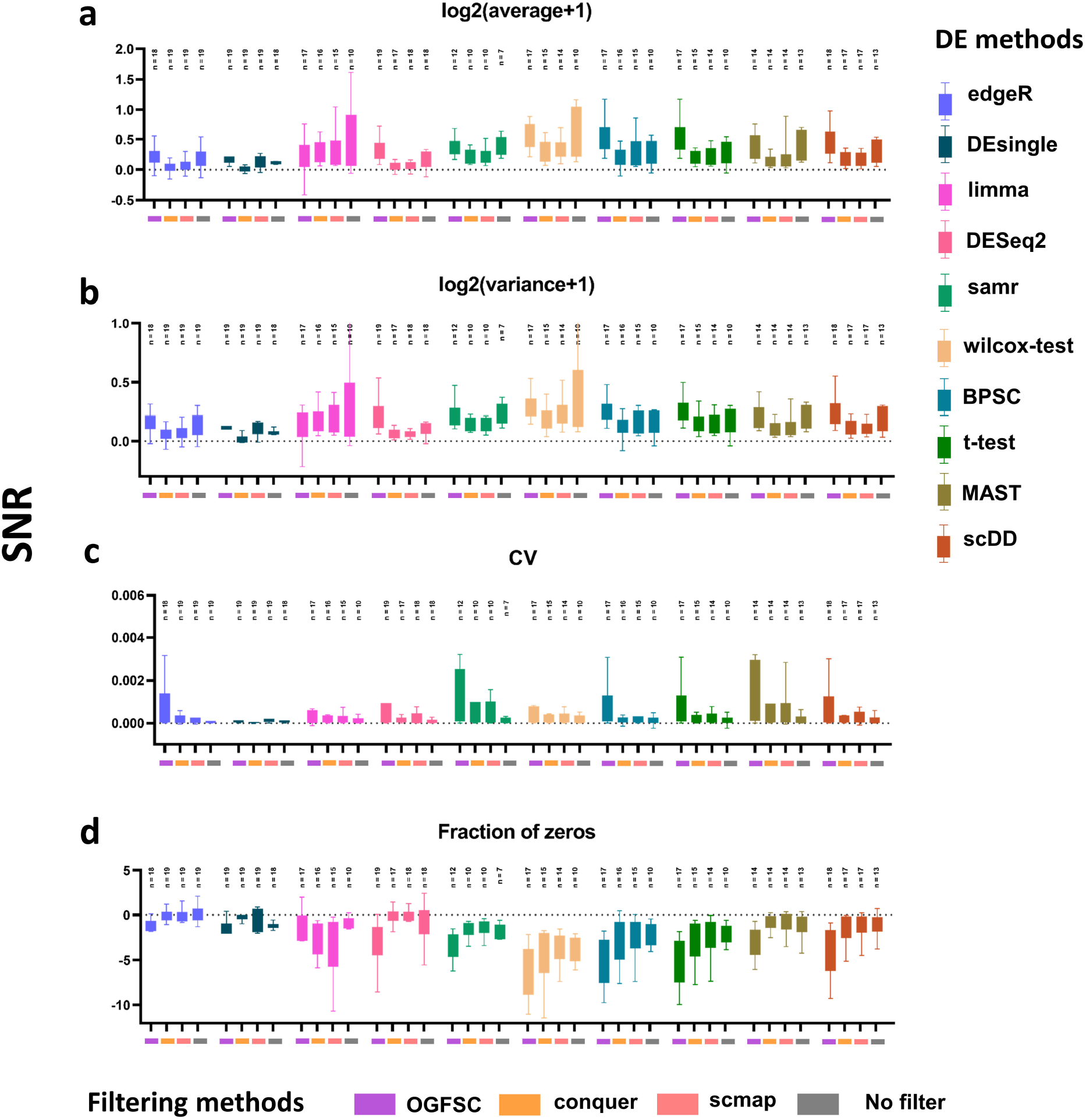
The SNR of DE gene detection on simulated datasets with true DE genes, exhibited by 4 characteristics: (a) SNR of log2(average+1). (b) SNR of log2(variance+1). (c) SNR of cv. (d) SNR of fraction of zeros. The color of box represents different DE methods; the colored rectangle represents different filtering methods. Dotted line, SNR = 0; n, number of simulated datasets. Outliers (larger than third quantile + 2*IQR or smaller than first quantile – 2*IQR) are excluded.

### The evaluation of FDP, TPR and AUROC on simulated data with true signal

Using the simulated data with a proportion of true DE genes, we investigated the performance of FDP, TPR and AUROC of different analysis strategies (Figure. 4). The FDP values of different methods varied in a wide range, some methods such as SAMseq, BPSC and scDD could keep a low FDP level in DE gene detection, but other methods including DESeq2 and limma, trended to result in a higher FDP (Figure. 4a). By comparing the performance of different filtering methods, we observed that all of the three filtering methods generally achieved lower FDP. For the DE methods which achieved low FDP like MAST and BPSC, false positive error could be alleviated by coupling with OGFSC. Filtering genes by conquer and scmap significantly improved the TPR value for all DE methods (Figure. 4b).

**Figure. 4.**
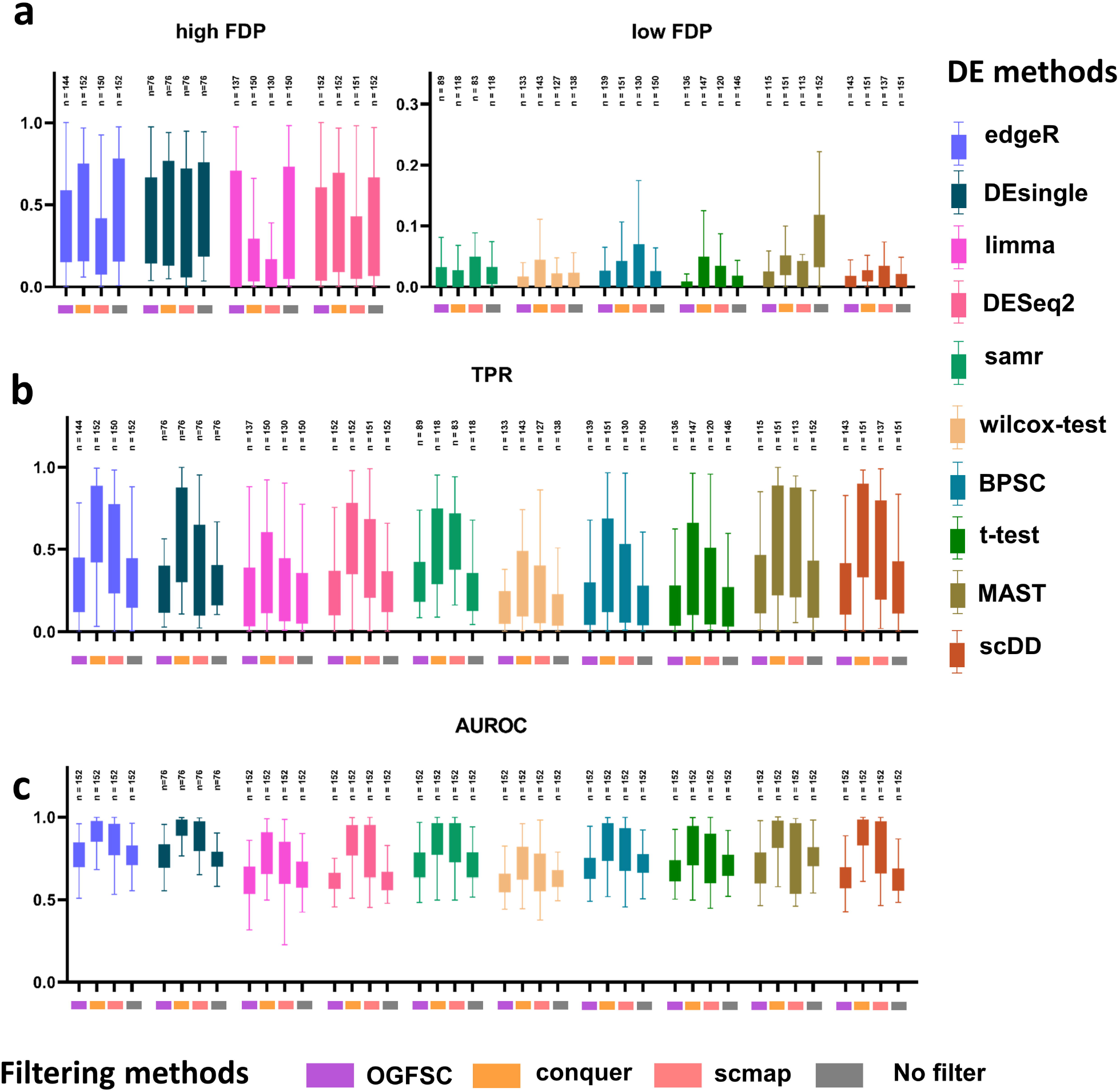
The Evaluation of FDP, TPR and AUROC on multiple simulated datasets. (a) Observed FDP with an adjusted p-value cutoff of 0.05, separated by high FDP (average > 0.2) and low FDP (average FDP < 0.1). (b) Observed TPR with an adjusted p-value cutoff of 0.05. (c) Observed AUROC with an adjusted p-value cutoff of 0.05. The colored boxes represent different DE methods; the lower horizontal-colored rectangles represent different filtering methods. n, number of simulated datasets. Outliers (larger than third quantile + 2*IQR or smaller than first quantile – 2*IQR) are excluded.

AUROC can be considered as a direct and gold-standard metric to assess the performance of methods. We observed that a lower FDP did not necessarily mean a higher AUROC, for example, although MAST and scDD achieve lower FDP than edgeR and limma, they had similar AUROC value (Figure. 4c). Gene filtering had a significant impact on AUROC values. The performances of DE gene detection with gene filtering method varied a lot. If no-filter was performed, the AUROC value would range between only 0.6 to 0.8. It’s remarkable that filtering genes by conquer and scmap could boost the AUROC up to 0.95 for DE methods like edgeR and scDD, and the average level for most DE methods would range between 0.8 to 0.9. The evaluation of TPR and AUROC indicated that the overall performances of most DE methods were generally similar, while filtering method seemed to be the one making the key difference (Figure. 4b, c).

### Metrics for personalized assessment in scCODE

The previous sections have demonstrated that filtering out genes containing much noise could be of great beneficial to DE gene detection in scRNA-seq data, therefore, the performance of DE gene detection depends on both gene filtering methods and DE methods. However, we found that the AUROC values for different datasets were divergent, which suggests that the appropriate analysis strategy for different dataset may differ (Figure. 5a). It implies there needs a personalized analyzing strategy for each specific scRNA-seq dataset. However, existing metrics, such as AUROC, which requires known true results, could not be used on real scRNA-seq dataset prior knowledge of truth DE genes. Thus, we developed CDO to imitate AUROC in real scRNA-seq datasets, and applied AUCC with the concept that consensus results could be more reliable.

**Figure. 5.**
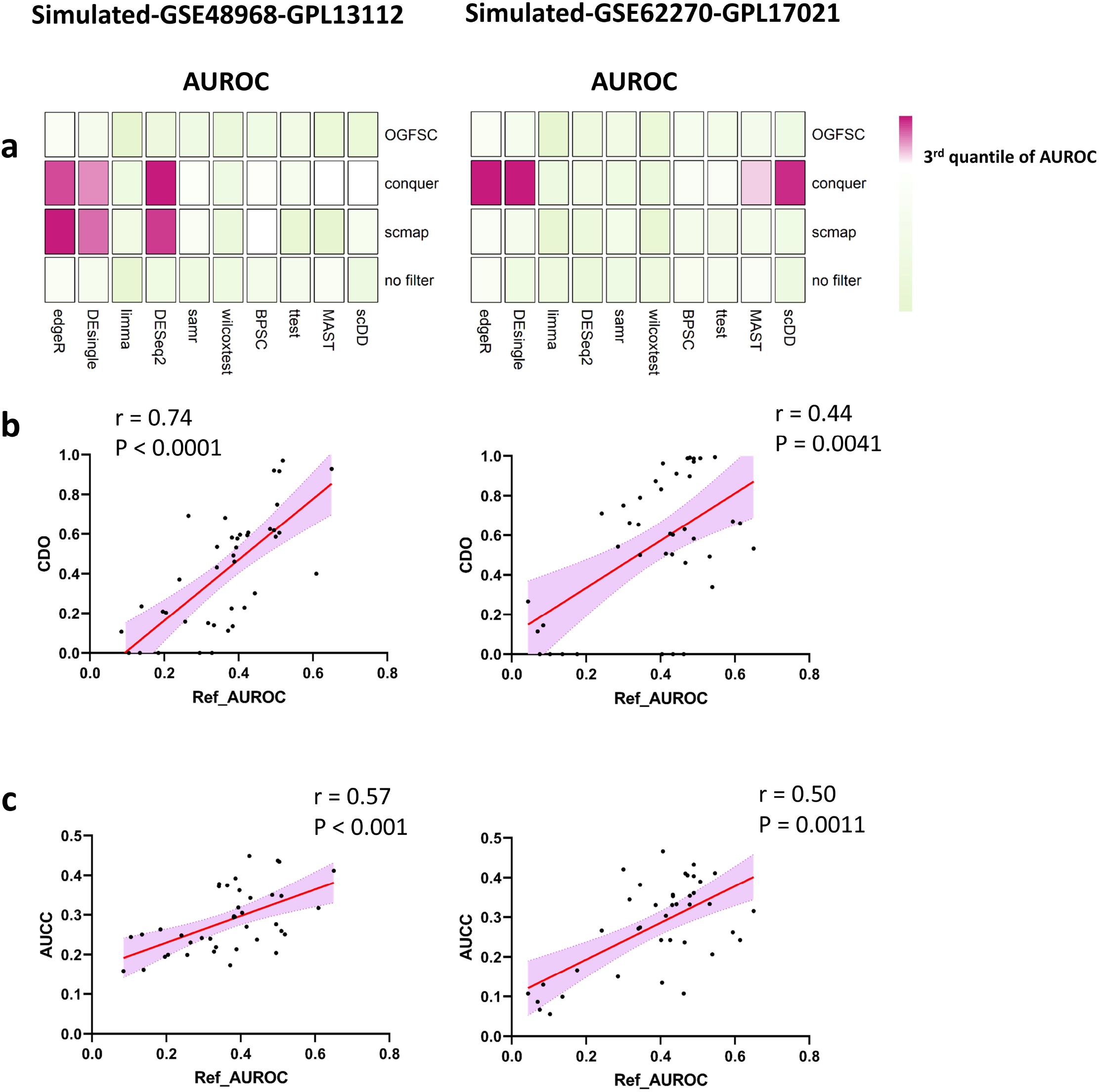
The reference AUROC of two simulated data with known true DE genes, and the spearman correlation between CDO, AUCC and AUROC. Left, Simulated-GSE48968-GPL13112 dataset; right, Simulated-GSE62270-GPL17021 dataset. (a) The heatmap of observed AUROC resulted by 40 analysis strategies on the two simulated data. (b) The correlation between CDO and AUROC, each point represents the result of one analysis strategy. (c) The correlation between AUCC and AUROC, each point represents the result of one analysis strategy.

We validated the performance of CDO and AUCC on 19 simulated datasets with 2% of genes randomly selected as true DE genes. We recorded the results of each dataset individually, and the corresponding AUROC results were used as reference. The “good” methods suggested by AUROC were largely divergent for different datasets (Figure. 5a). We further investigated the correlation between CDO, AUCC and AUROC for each simulated dataset. For most datasets, the correlation between CDO and the reference AUROC was significant (Figure. 5b, Supplementary figure. 3). The AUCC was also highly correlated with AUROC for most simulated datasets (Figure. 5C, Supplementary figure. 4). Consistent to the mathematic derivation presented in the Methods section, we here demonstrate CDO and AUCC are reliable approximates of AUROC in real scenarios, and therefore we propose to use the two metrics in scCODE, which makes it practical for personalized analysis strategy.

### scCODE benchmarked on simulated datasets

Based on the metrics CDO and AUCC, scCODE could access the performance of different analysis strategies on experimental scRNA-seq datasets. As mentioned before, we consider the DE genes detected by majority of analysis strategies as more reliable, and make a further consensus optimization of the DE gene results detected by the “suitable” analysis strategies assigned by scCODE. Finally, scCODE provides the consensus optimized DE gene results as DE gene output.

Here, we benchmarked the performance of scCODE against other analysis strategies on several simulated datasets. The sensitivity and specificity of the DE results were considered, a higher TPR of the top DE genes would indicate better performance. The same analysis strategy applied to different datasets obtained divergent performances. scCODE achieved higher TPR than most analysis strategies, which indicated that scCODE could identify DE genes more accurately for these simulated datasets (Figure. 6). Besides, the performance of scCODE was more stable on different simulated datasets.

**Figure. 6.**
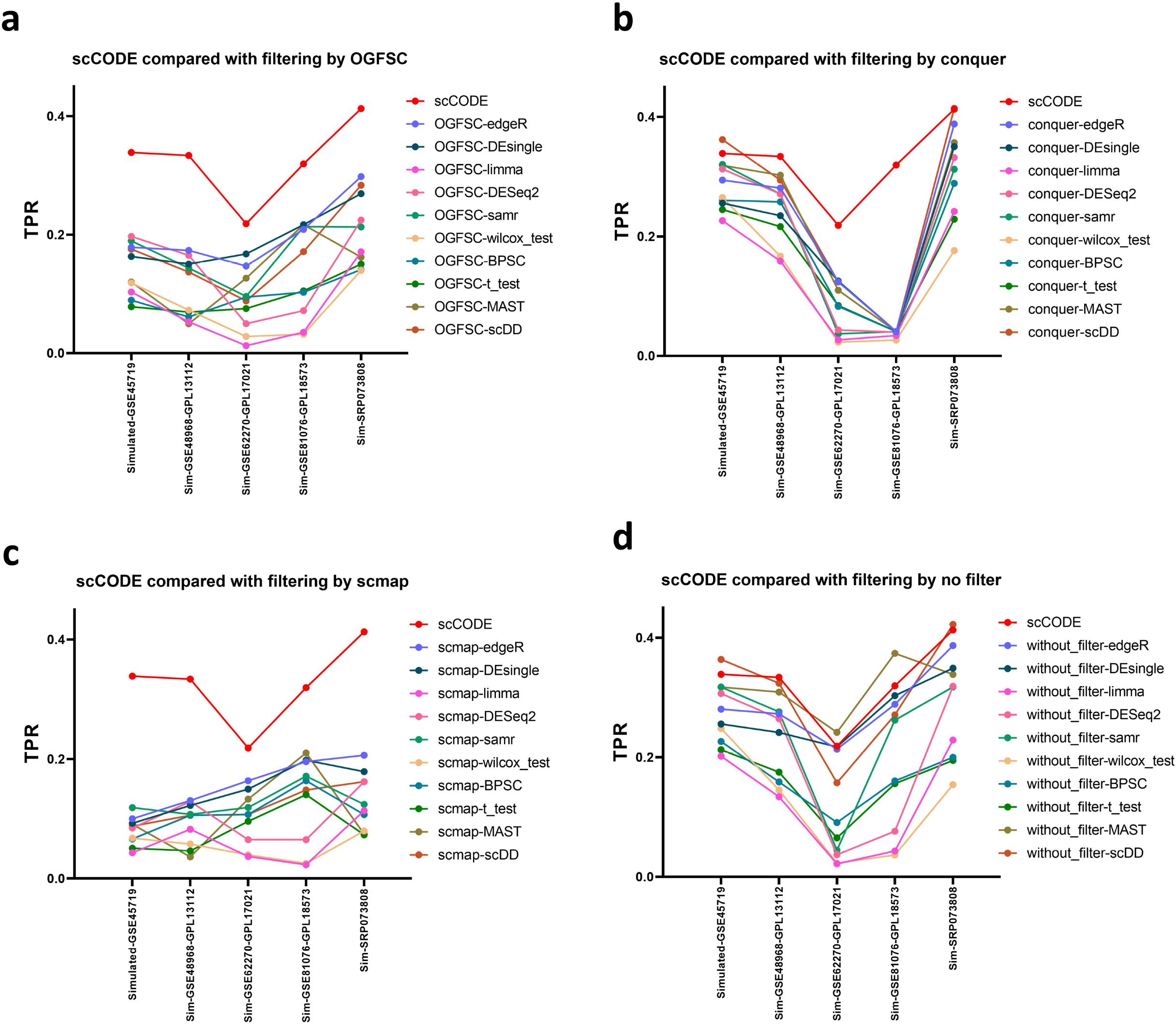
scCODE benchmarked on simulated datasets. (a) scCODE compared with the analysis strategies filtering by OGFSC, the TPR was calculated on the top n DE gene results detected by each analysis strategy, where n = the total number of DE genes set in each simulated dataset. (b) scCODE compared with analysis strategies filtering by conquer. (c) scCODE compared with analysis strategies filtering by scmap. (d) scCODE compared with analysis strategies without filtering genes.

### scCODE validated on a real dataset

In this article, we also applied scCODE on a real scRNA-seq dataset from immune-stimulation research of mouse[19], containing stimulated CD4+ T cells (by antibodies to CD3 ε and CD28) and naive ones (Figure. 7). The heatmap represented the normalized results of CDO and AUCC, which exhibited the appropriateness of various analysis strategy for the data. We selected 5 suitable analysis strategies (i.e., OGFSC-Wilcoxon-test, conquer-limma, etc.), and 5 unsuitable analysis strategies (i.e., OGFSC-MAST, no-filter-DEsingle, etc.) (Figure. 7a). Then, for each strategy, top DE genes with the smallest p-values (all with p<0.05) were identified as candidate DE genes. The results from suitable analysis strategies were compared to the unsuitable ones. The suitable analysis strategies detected a total of 1055 unique DE genes, while the unsuitable ones detected 1546 unique genes (Figure. 7b). Besides, the suitable analysis strategies had more overlapped genes (more than 50% genes were detected ≥ 3 times) than that of unsuitable ones (only 1.3% genes were detected ≥ 3 times). It indicated that the suitable strategies trend to achieve more consistent results.

**Figure. 7.**
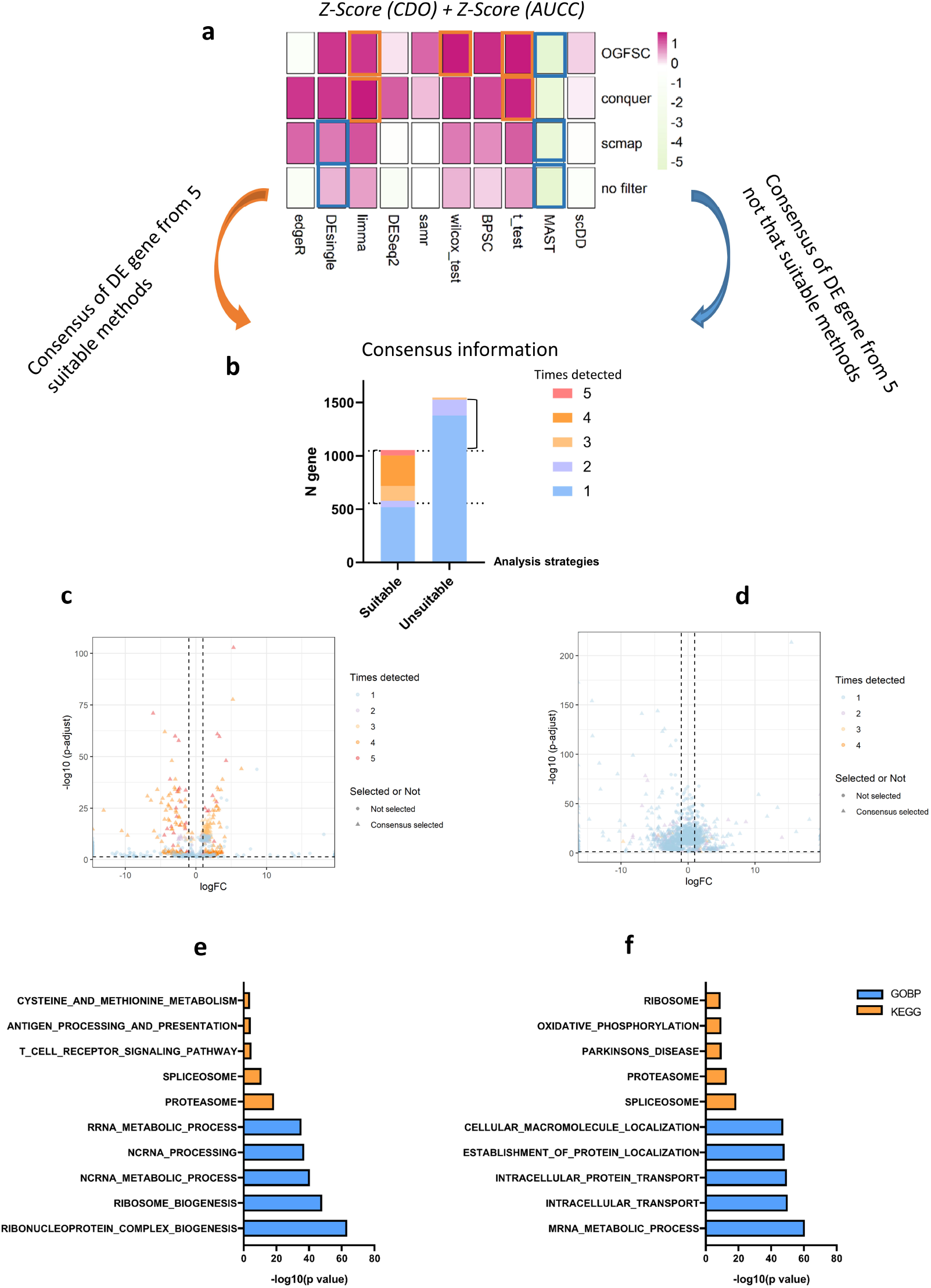
scCODE validated on a real scRNA-seq data comparing activated CD4+ T cells with naïve ones. (a) Heatmap of the average results of CDO and AUCC. White rectangle, 5 suitable analysis strategies suggested by scCODE; black rectangle, top 5 unsuitable analysis strategies. (b) Bar plot of the DE gene information of the 5 suitable and unsuitable analysis strategies. Dotted line, consensus selection line, the genes above the line were selected. (c, d) Volcano map and consensus information of the top 500 DE genes collected from each of the analysis strategy; (e) suitable ((f) unsuitable) analysis strategies. Dotted line, p-adjusted = 0.05, logFC = ± 1. Detected times, gene detected times within the 5 analysis strategies selected. (e, f) GOBP and KEGG analyze of the up-regulated genes in activated CD4+ T cells from consensus DE genes of (e) suitable and (f) unsuitable analysis strategies. Top 5 results of GOBP and KEGG are exhibited.

Furthermore, we also compared the results after consensus ranking and selection (Figure. 7 c, d). The up-regulated genes of stimulated CD4+ T cells were processed by gene set enrichment analysis (Gene Ontology (GO)-biological process (BP) and KEGG pathway), and top 5 results were exhibited (Figure. 7e, f). We found that all of the enriched GOBP functions uncovered by suitable analysis strategies, such as RIBOSOME_BIOGENESIS, NCRNA_PROCESSING and RRNA_METABOLIC_PROCESS, were in profound agreement with the conclusion of the original article[19]; while that was not the case for the unsuitable results. And the KEGG results of suitable analysis strategies also detected a pathway (T_CELL_RECEPTOR_SIGNALING_PATHWAY) closely associated with T cell activation. The consensual DE gene results from the suitable analysis strategies of methods given by scCODE were more closely matched with the experimental question studied.

## Discussion

DE gene detection in scRNA-seq data is one of the most concerned steps in data analyses. Gene filtering is generally applied before DE gene detection, but the influence of gene filtering is not fully recognized before. In this article, we first presented a systematic benchmark on both gene filtering methods and DE methods for scRNA-seq data. We exhibited the results from the aspects of FPR, FDP, SNR and AUROC. Without considering the influence of gene filtering, we got results in agreement with previous researches[15, 16]. Some methods like MAST and Wilcoxon-test performed well in specificity, and trended to detect less false DE genes (low FPR and FDP). But not a single DE method showed a superiority in sensitivity (TPR), and most DE methods achieved similar performance by AUROC. The performances with gene filtering methods were divergent, and the improvement it brought was significant. We have shown that gene filtering does help DE gene detection in all aspects. We also summarized the evaluation on multiple simulated datasets (Supplementary figure. 5). The summary categorized each analysis strategy into “Good”, “intermediate” and “Poor” on each metric. It also showed that the performance of DE gene detection depended on both gene filtering and DE processing.

Previous evaluation tools were mainly based on evaluating performance with cohort of datasets. We have shown that the performance of DE gene detection on different datasets varies, and personalized analytical strategy on the specific dataset studied is necessary. To assess different methods in practical DE detection studies, two novel metrics CDO and AUCC were proposed in this study. Theoretically, the new metrics could faithfully assess a DE gene detection result; and it was also approved on simulated datasets by demonstrating significant correlation between the proposed metrics and AUROC as reference. In fact, the reliability of CDO is based on the hypothesis that the seed genes are true DE ones. We considered the common DE genes detected by different analysis strategies as seed genes, and investigated the threshold to guarantee its reliability. We validated the relationship between the threshold of seed gene selection (genes detected in how many percent of strategies) and significancy of correlation between CDO and AUROC (Supplementary figure. 6). The results on simulated data created by splatter[28] showed that the reliability of CDO was highly dependent on the threshold (Supplementary figure. 6). A low threshold tended to result in high false-positive rate in seed gene selection, while a too high threshold would sharply reduce the number of seed genes and its stability. Thus, we suggested and applied a threshold of seed genes higher than 75%, and restricted the number of seed genes no less than 5 in this article.

The R package based on the two metrics, scCODE, answered the question of which analysis strategies are appropriate for a scRNA-seq dataset. More importantly, it can provide optimized DE gene detection results of experimental datasets. We validated scCODE on both simulated datasets and a real scRNA-seq dataset. Figure. 6 showed that scCODE achieved better and more robust results than most analysis strategies on each dataset, as it judged and applied personalized optimal strategies on these datasets, while the other analysis strategies performed unstably on different datasets. Besides, the validation on the experimental scRNA-seq dataset exhibited that scCODE could provide reliable assessment of these strategies on real scRNA-seq dataset, and could further provide reliable DE gene results for the users.

scCODE currently contains 4 filtering methods and 10 DE methods, it automatically produces the optimal DE gene results for the given dataset (A demo output of scCODE could be seen in Supplementary table. 2). A limitation of scCODE is that it needs a comparison of multiple methods to find the seed DE genes, thus it may be computationally demanding. We also provided a light version of scCODE including only a few reliable methods, such as scDD, MAST and DESeq2 (Supplementary figure. 5a). The light version provided very similar result on the experimental data tested (Supplementary figure. 5b). We believe that scCODE is convenient to conduct optimized DE gene detection for specific scRNA-seq data. In the future, more methods will be incorporated in scCODE. Besides, there may be other preprocessing steps which may be included in the evaluation framework of scCODE, such as normalization[29, 30], etc.

## Conclusion

In this study, we systematically evaluated the true influence of gene filtering on DE gene detection. The performance of DE gene detection depends on both gene filtering and DE processing. And we found that personalized analysis strategy is needed to achieve optimal DE gene results for each experimental scRNA-seq dataset. However, existing metrics requiring prior truth DE genes were not suitable for experimental datasets. We established two novel metrics CDO and AUCC for it, and observed significant correlation between the new metrics and AUROC (regarded as ground truth) on multiple simulated datasets independently. Finally, we developed scCODE, a R package for optimizing DE gene detection of scRNA-seq dataset. scCODE performed better and more robust on different simulated datasets. The consensus optimized DE gene results suggested by scCODE better match the experimental design of the test data. We hope our user-friendly tool scCODE could help the analysis of each scRNA-seq dataset, and could be beneficial to maintain more reliable DE gene results.

## Declarations

### Ethics approval and consent to participate

Not applicable.

### Consent for publication

Not applicable.

### Availability of data and materials

All of the code were based on R version 4.02.

The origin data used in evaluation are available in the conquer repository, at http://imlspenticton.uzh.ch:3838/conquer/.

The experimental data used to test scCODE is available in the reference paper[19], and also can be found at https://github.com/XZouProjects/scCODE/tree/main/data.

The R code for evaluation of gene filtering and DE methods is available at https://github.com/XZouProjects/scCODE/evaluation.

The installable version of the scCODE R package is available at https://github.com/XZouProjects/scCODE.

### Authors’ contributions

JWZ, JH, XZ, KC conceived and designed the algorithm. JWZ developed the software. JH, XZ, KC, MCW, ZZ and ZQL provided data interpretation and biological explanation. JWZ, JH, XZ and KC wrote the manuscript, JH, XZ, KC, JWZ and MCW prepared figures. RH and XBZ provided critical suggestions. The manuscript was approved by all authors.

## Acknowledgements

We thank Li Xuejing for her help of Figure. 2.

The data analysis was supported by GorgeousAI (Gu Ai) Co. Ltd.

## Additional files

1. Supplementary table. 1 A excel lists the detail information of the experimental datasets and the simulated datasets, which were used for evaluation of DE gene detection performance.

2. Supplementary table. 2. A excel lists the demo results of scCODE applied on the immune-senescence dataset in this article. It contains 4 sheets, the results of Z-Score of AUCC, the results of Z-Score of CDO, the strategy information selected (default as top 5) and the information of the final DE genes results.

3. Supplementary figures. A PPT file includes all the supplementary figures in sequence. The figure legend and description are list in PPT comment of each page.

