## Supplementary figures for "scCODE: an R package for personalized differentially expressed gene detection on single-cell RNA-sequencing data"

### Slide 1
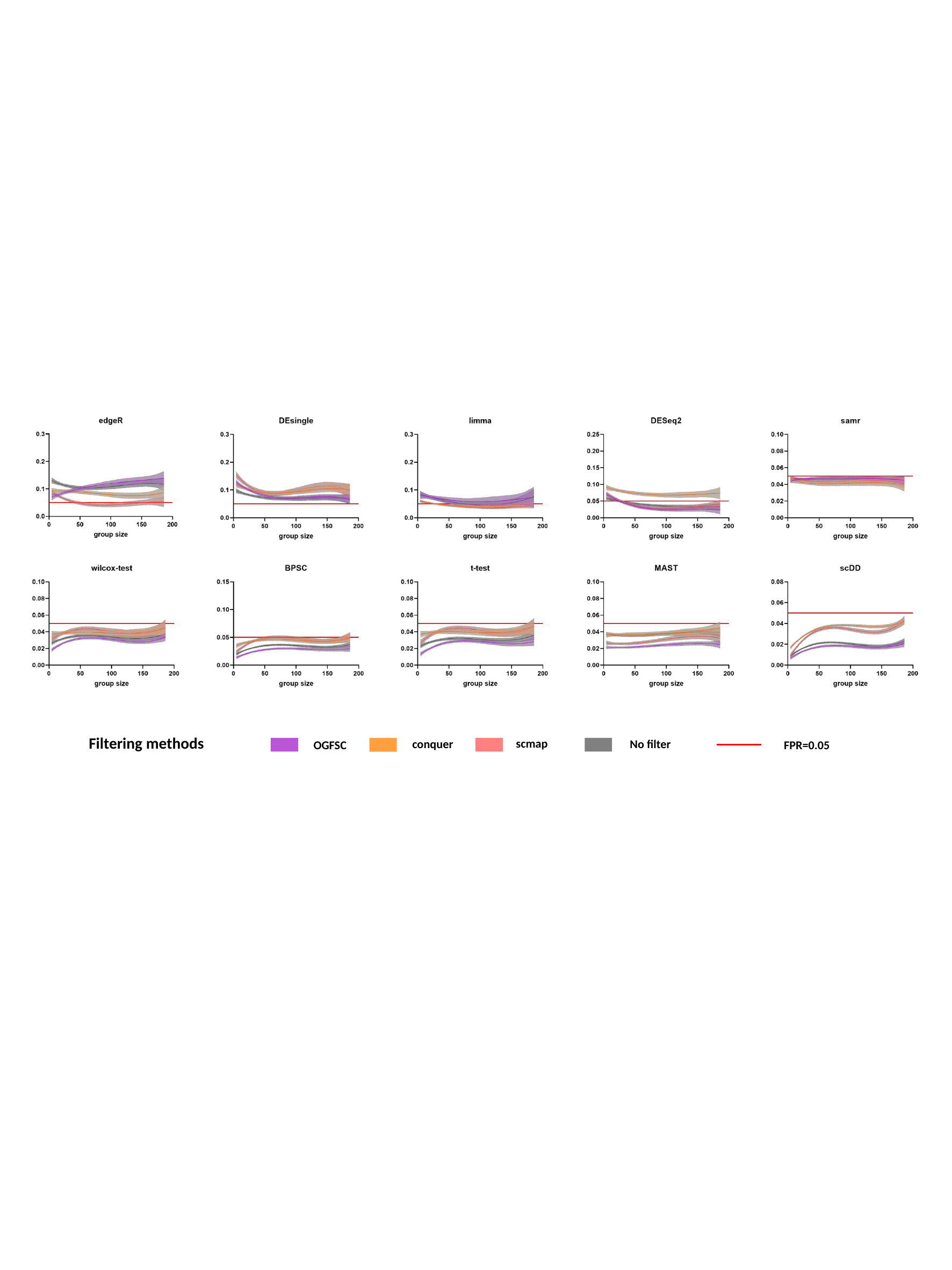

Filtering methods
scmap
conquer
No filter
OGFSC
FPR=0.05

### Slide 2
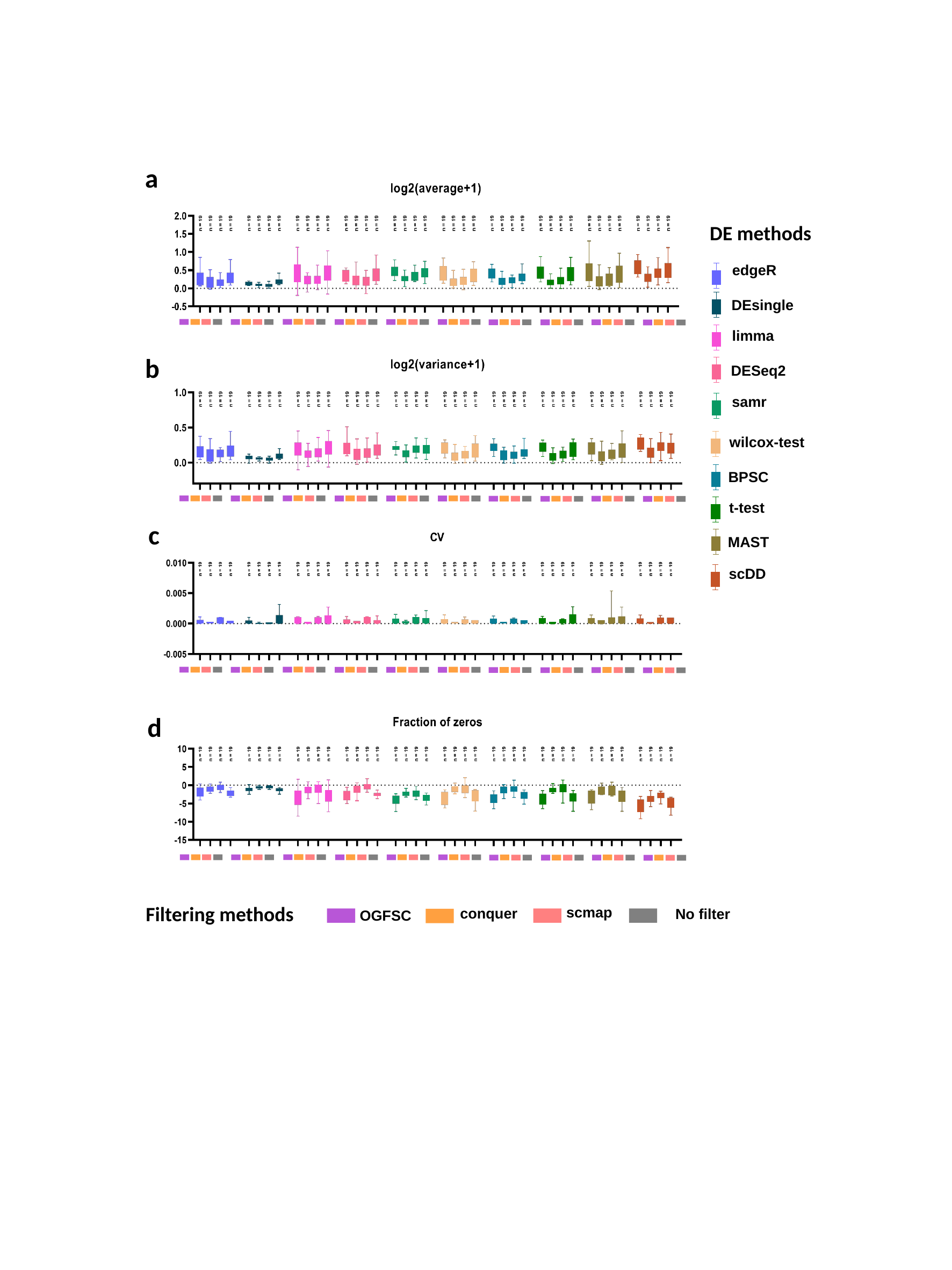

a
DE methods
edgeR
DEsingle
limma
DESeq2
samr
wilcox-test
BPSC
t-test
MAST
scDD
b
c
d
Filtering methods
scmap
conquer
No filter
OGFSC

### Slide 3
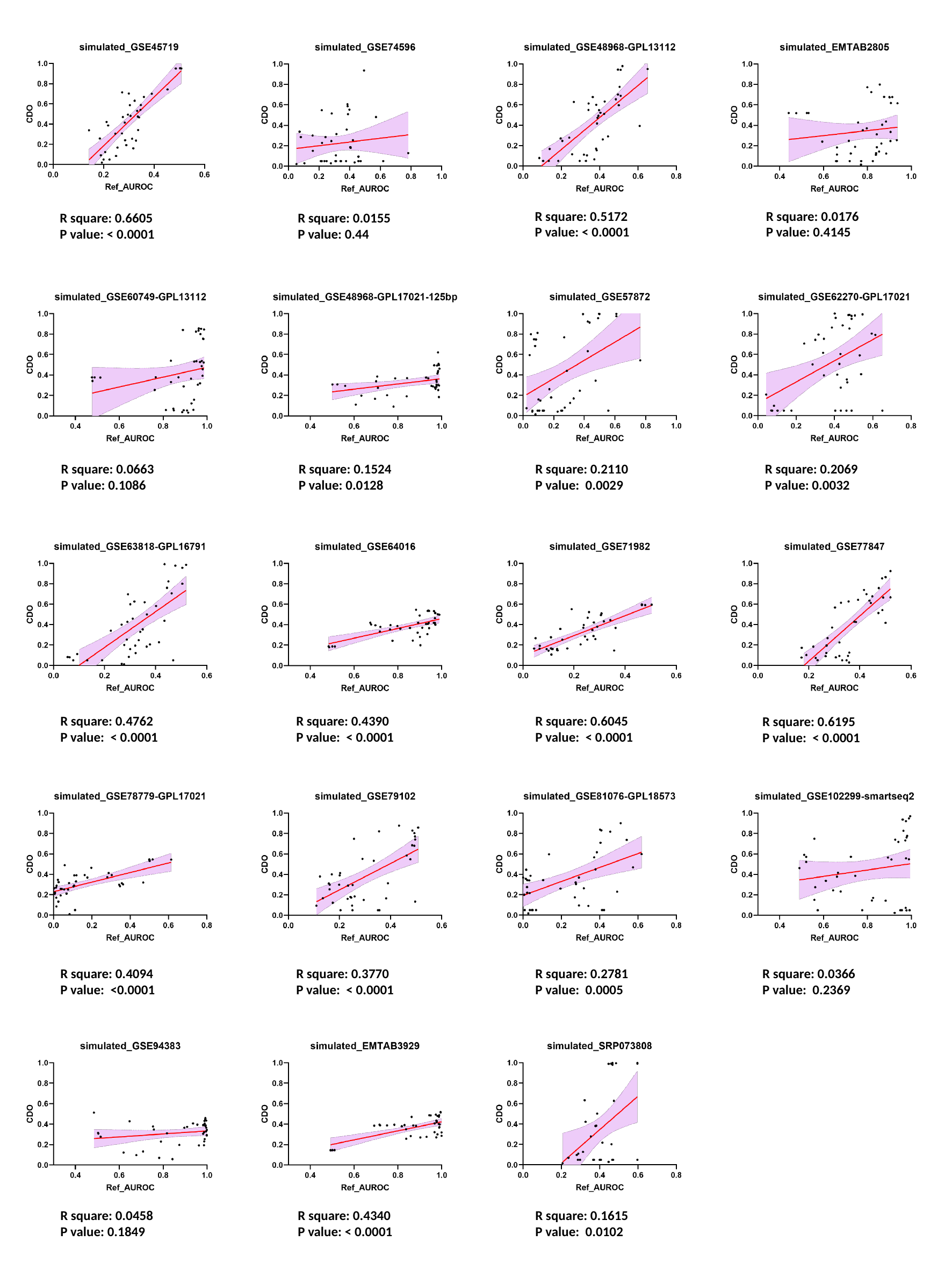

R square: 0.5172
P value: < 0.0001
R square: 0.0176
P value: 0.4145
R square: 0.0155
P value: 0.44
R square: 0.6605
P value: < 0.0001
R square: 0.1524
P value: 0.0128
R square: 0.2110
P value: 0.0029
R square: 0.2069
P value: 0.0032
R square: 0.0663
P value: 0.1086
R square: 0.4390
P value: < 0.0001
R square: 0.6045
P value: < 0.0001
R square: 0.4762
P value: < 0.0001
R square: 0.6195
P value: < 0.0001
R square: 0.3770
P value: < 0.0001
R square: 0.2781
P value: 0.0005
R square: 0.4094
P value: <0.0001
R square: 0.0366
P value: 0.2369
R square: 0.4340
P value: < 0.0001
R square: 0.1615
P value: 0.0102
R square: 0.0458
P value: 0.1849

### Slide 4
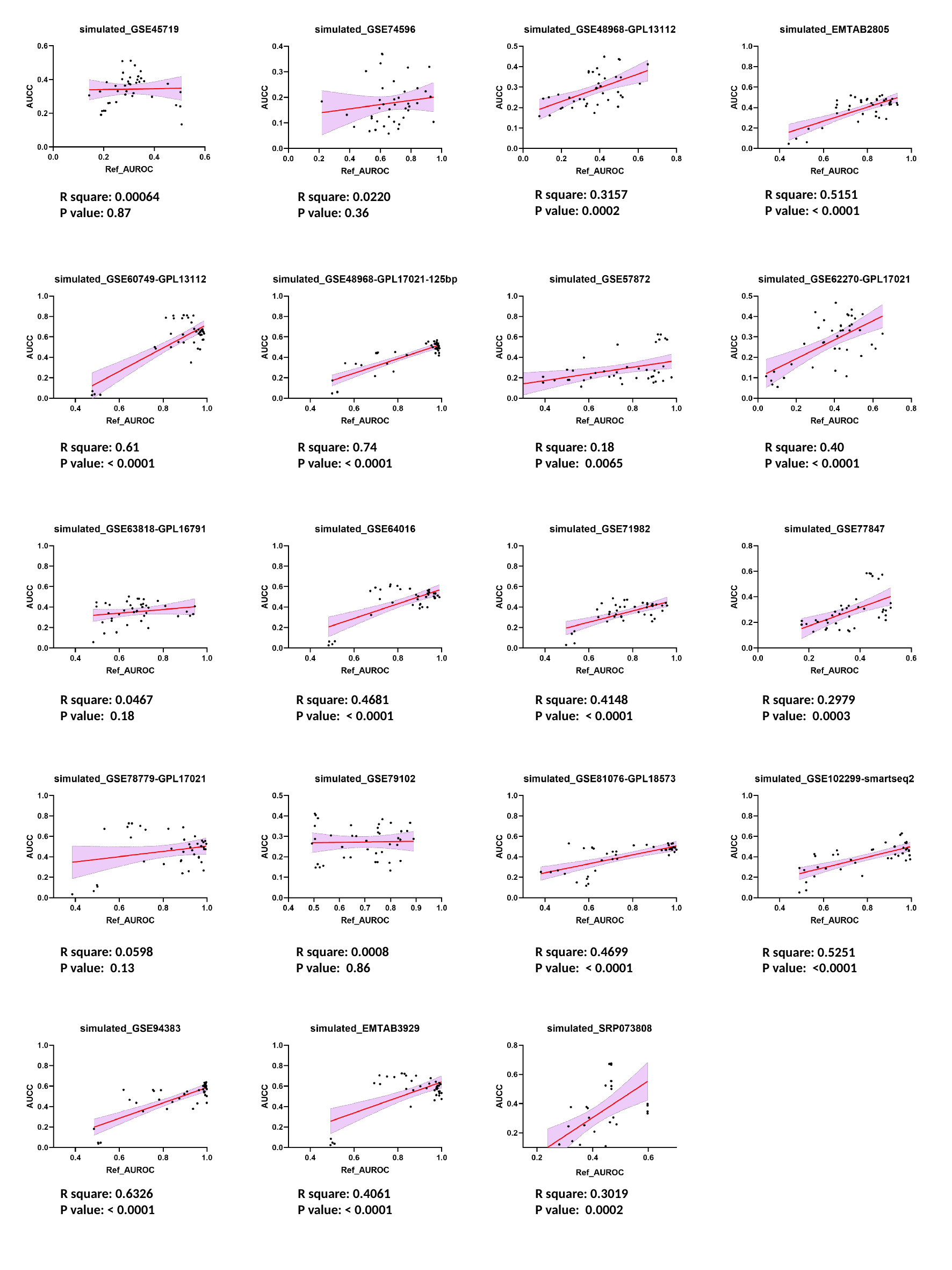

R square: 0.3157
P value: 0.0002
R square: 0.5151
P value: < 0.0001
R square: 0.0220
P value: 0.36
R square: 0.00064
P value: 0.87
R square: 0.74
P value: < 0.0001
R square: 0.18
P value: 0.0065
R square: 0.40
P value: < 0.0001
R square: 0.61
P value: < 0.0001
R square: 0.4681
P value: < 0.0001
R square: 0.4148
P value: < 0.0001
R square: 0.0467
P value: 0.18
R square: 0.2979
P value: 0.0003
R square: 0.0008
P value: 0.86
R square: 0.4699
P value: < 0.0001
R square: 0.0598
P value: 0.13
R square: 0.5251
P value: <0.0001
R square: 0.4061
P value: < 0.0001
R square: 0.3019
P value: 0.0002
R square: 0.6326
P value: < 0.0001

### Slide 5
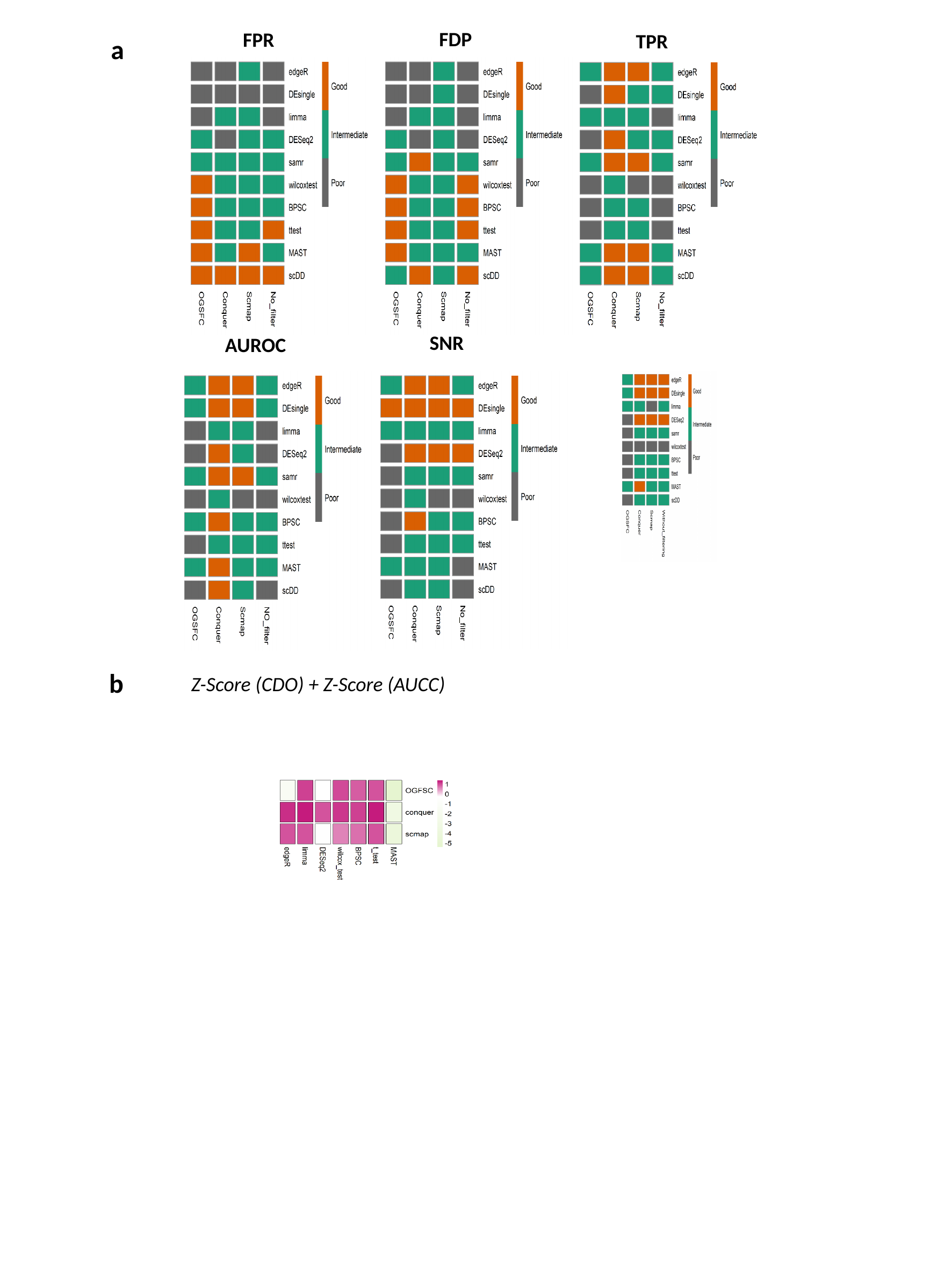

FDP
FPR
TPR
SNR
AUROC
a
b
Z-Score (CDO) + Z-Score (AUCC)

### Slide 6
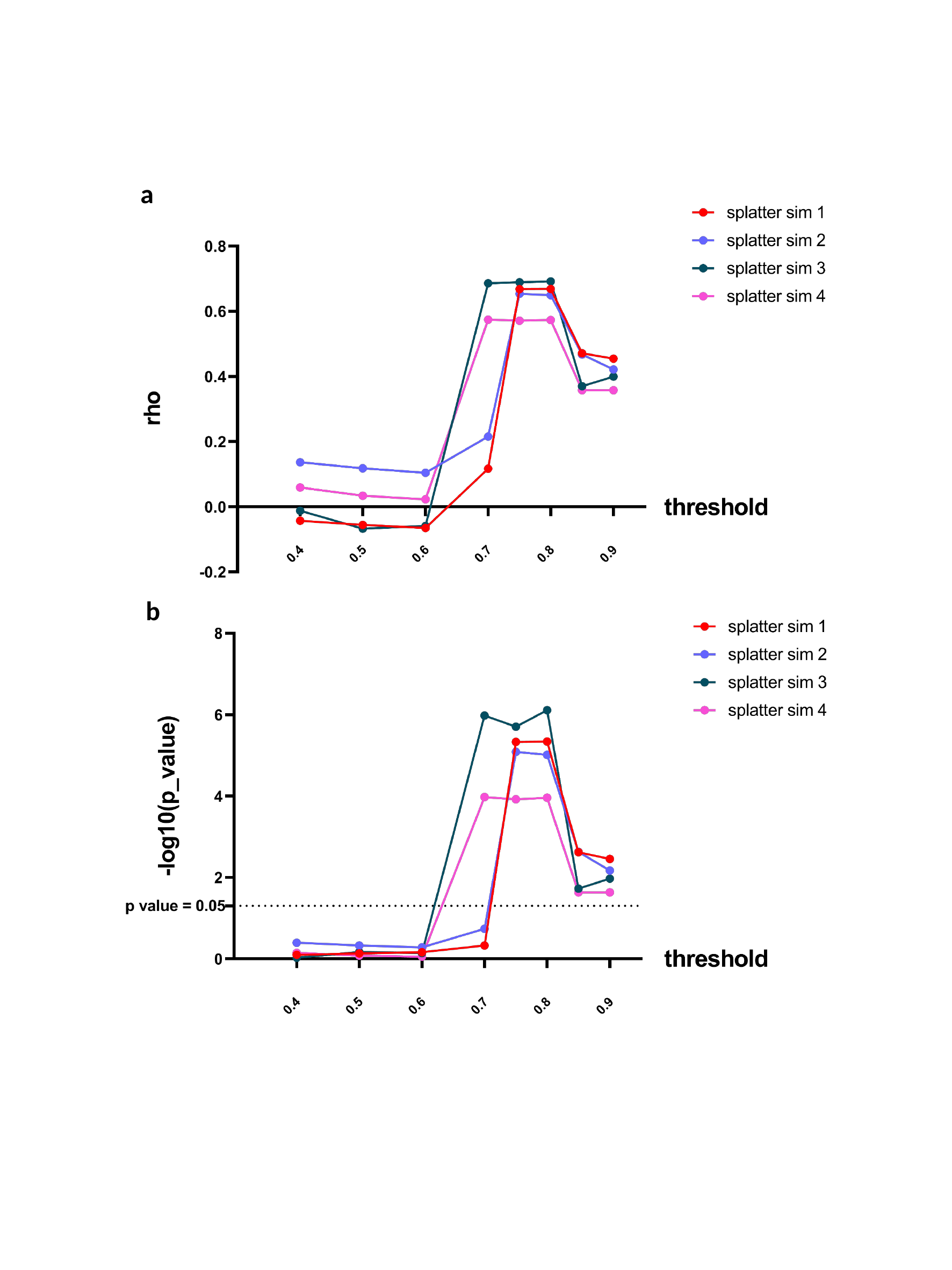

a
b
